# The genomes of precision edited cloned calves show no evidence for off-target events or increased *de novo* mutagenesis

**DOI:** 10.1101/2021.01.28.428703

**Authors:** Swati Jivanji, Chad Harland, Sally Cole, Brigid Brophy, Dorian Garrick, Russell Snell, Mathew Littlejohn, Götz Laible

## Abstract

Animal health and welfare are at the forefront of public concern and the agricultural sector is responding by prioritising the selection of welfare-relevant traits in their breeding schemes. In some cases, welfare-enhancing traits such as horn-status (i.e., polled) or diluted coat colour, which could enhance heat tolerance, may not segregate in breeds of primary interest, highlighting gene-editing tools such as the CRISPR-Cas9 technology as an approach to rapidly introduce variation into these populations. A major limitation preventing the acceptance of CRISPR-Cas9 mediated gene-editing, however, is the potential for off-target mutagenesis, which has raised concerns about the safety and ultimate applicability of this technology. Here, we present a clone-based study design that has allowed a detailed investigation of off-target and *de novo* mutagenesis in a cattle line bearing edits in the *PMEL* gene for diluted coat-colour. No off-target events were detected from high depth whole genome sequencing performed in precursor cell-lines and resultant calves cloned from those edited and non-edited cell lines. Long molecule sequencing at the edited site and plasmid-specific PCRs did not reveal structural variations and/or plasmid integration events in edited samples. Furthermore, an in-depth analysis of *de novo* mutations across samples revealed that the mutation frequency and spectra were unaffected by editing status. Cells in culture, however, had a distinct mutation signature where *de novo* mutations were predominantly C>A mutations, and in cloned calves they were predominantly T>G mutations, deviating from the expected excess of C>T mutations. We conclude that the gene-edited cells and calves in this study did not present a higher mutation load than unedited controls. Cell culture and somatic cell nuclear transfer cloning processes contributed the major source of contrast in mutational profile between samples.

## Introduction

The agriculture sector’s response to demands for enhanced animal welfare, production, efficiency and sustainability is sometimes limited by available genetic variation within a particular population. Although favourable variation may be introgressed from other populations by cross-breeding, stabilising favourable variation by selective breeding regimes typically comes at the cost of losses in genetic gain and inbreeding depression. Gene-editing offers an attractive solution with its ability to directly introduce precise polymorphisms causal for favourable traits within a single generation. Acceptance of gene editing technologies is in part dependent on the occurrence of mutagenesis at sites other than the intended on-target site, or ‘off-target’ mutagenesis, and the ability to detect these events above baseline mutation levels. The clustered regularly interspaced short palindromic repeat (CRISPR)-CRISPR associated (Cas) system is a versatile and popular gene-editing tool proven to be successful in large animal models (1). The most commonly used CRISPR-Cas9 system is derived from *Streptococcus pyogenes*, and uses the Cas9 endonuclease complexed with a guide RNA (gRNA) that identifies and binds to a 20 nucleotide target region (protospacer) immediately preceding a NGG protospacer-associated motif, or PAM. The endonuclease induces a double stranded break 3bp upstream of the NGG site, which can either be repaired via non-homologous end joining, or a repair template coding for the desired polymorphism can be introduced to facilitate homology-directed repair (2,3). The potential for off-target mutations have been associated with non-unique matches and sequence mismatches distal from the PAM sequences at the 5’ end of the gRNA (4–6). Structural variation at the targeted edit site (7–9), and unintended integration of the editing vectors (10,11) have also been associated with gene-editing and have raised concerns about the safety and applicability of these technologies in biomedicine and agriculture.

Off-target mutations have been investigated by amplification and sequencing of pre-selected sites identified by bioinformatic tools that highlight sequences with homology to the on-target site (12–14). This method may not be practical for large-scale screening, with the generation of a large number of possible non-unique matches. This approach also neglects to consider the potential for mutations to be introduced at sites with low on-target sequence similarity, and thus will not be able to identify such events. Whole genome sequencing (WGS) is a less biased approach to off-target mutation detection and enables analysis of single nucleotide variants (SNV), small insertions and deletions (indels), and some structural variants (SV), that may arise as a result of the use of CRISPR-Cas9 mediated gene-editing. However, since cells naturally accumulate *de novo* mutations through spontaneous mutagenesis during cell division, it is challenging to distinguish mutations attributable to the application of gene-editing technologies from those that occur spontaneously. To characterise any off-target mutagenesis, one approach is to quantify changes in detectable *de novo* mutation between gene-edited samples and controls, and then assess whether candidate variants do, or do not, sit in biologically plausible off-target sites. This approach has been used to evaluate the presence and frequency of off-target mutations in gene-edited large animal models, generated from multiplexed single-cell-embryo injection, and their progeny (7,15). Wang et al. (7) and Li et al. (15) used a trio-based study design to investigate off-target effects of CRISPR-Cas9 and showed that the off-target mutation rate was negligible and the *de novo* mutation rate in edited animals was comparable to their non-edited controls. A WGS approach to off-target mutation detection was also used by Schaefer et al. (16) to identify off-target mutations in two gene-edited mice generated by single-cell embryo injection (17). Schaefer et al. (16) reported hundreds of off-target mutations by WGS comparison to a single untreated mouse, but this result was found to be flawed when the authors later reported no excess mutations upon conducting WGS analysis with additional mouse lines (18). These studies highlight the importance of considering inherited and spontaneous mutations when investigating off-target events, and the use of appropriate controls that enable these considerations to be made.

In this study, we conduct the first WGS analysis in cloned cattle generated from a gene-edited cell line to evaluate off-target events and *de novo* mutagenesis associated with the application of CRISPR-Cas9 mediated gene-editing and cloning to create live cattle for use in agriculture. We analysed WGS from a cell clone homozygous for a CRISPR-Cas9 induced 3bp deletion in the premelanosomal protein gene (*PMEL*), the parental (non-edited) primary fetal cell line that cell clone was derived from, as well as two edited and three control calves generated from these cells by somatic cell nuclear transfer. The 3bp deletion in the *PMEL* gene was proposed to cause coat colour dilution in Highland and Galloway cattle (19), and by introducing it into a Holstein-Friesian background, Laible et al. (20) simultaneously demonstrated causality of this mutation and introduced a favourable trait within a single generation (20). Taking advantage of the clone-based study design, we used WGS and other molecular approaches to comprehensively screen for off-target SNVs, indels, and SVs that could be attributed to the use of CRISPR-Cas9 mediated gene-editing. We found no detectable CRISPR-Cas9 associated off-target mutations, and that the *de novo* mutation rate in calves generated from the gene-edited cell line was no different in calves generated from the non-edited cell line of same parental origin.

## Results

### Origin of the study material and analysis of whole genome sequence data

We used the recently described cloned calves that were edited for a *PMEL* coat colour dilution mutation (20) to investigate the precision of CRISPR-Cas9 gene-editing. For our in-depth genotype analysis, we applied WGS and included the parental, non-edited cell line (BEF2), the gene-edited clonal cell line (CC14) derived from BEF2, three control clones (1802, 1803 and 1804) generated from BEF2 cells, and two gene-edited clones (1805 and B071) that were generated with CC14 donor cells (Fig 1). The average whole genome sequencing depth per sample was 50.7x, after alignment to the bovine reference genome ARS-UCD1.2 (21). Greater than 99% of the reads mapped to the reference genome, and more than 92% of the reads mapped with a map quality score of 60 across all samples except sample B071, which had approximately 80% of reads with a map quality score of 60. Variant calling using GATK HaplotypeCaller (22) identified 8,021,969 variants across the seven samples. A pair-wise genomic concordance test across the seven samples found 99.99% concordance between all pairs, consistent with clones originating from the same genetic background.

**Fig 1.**
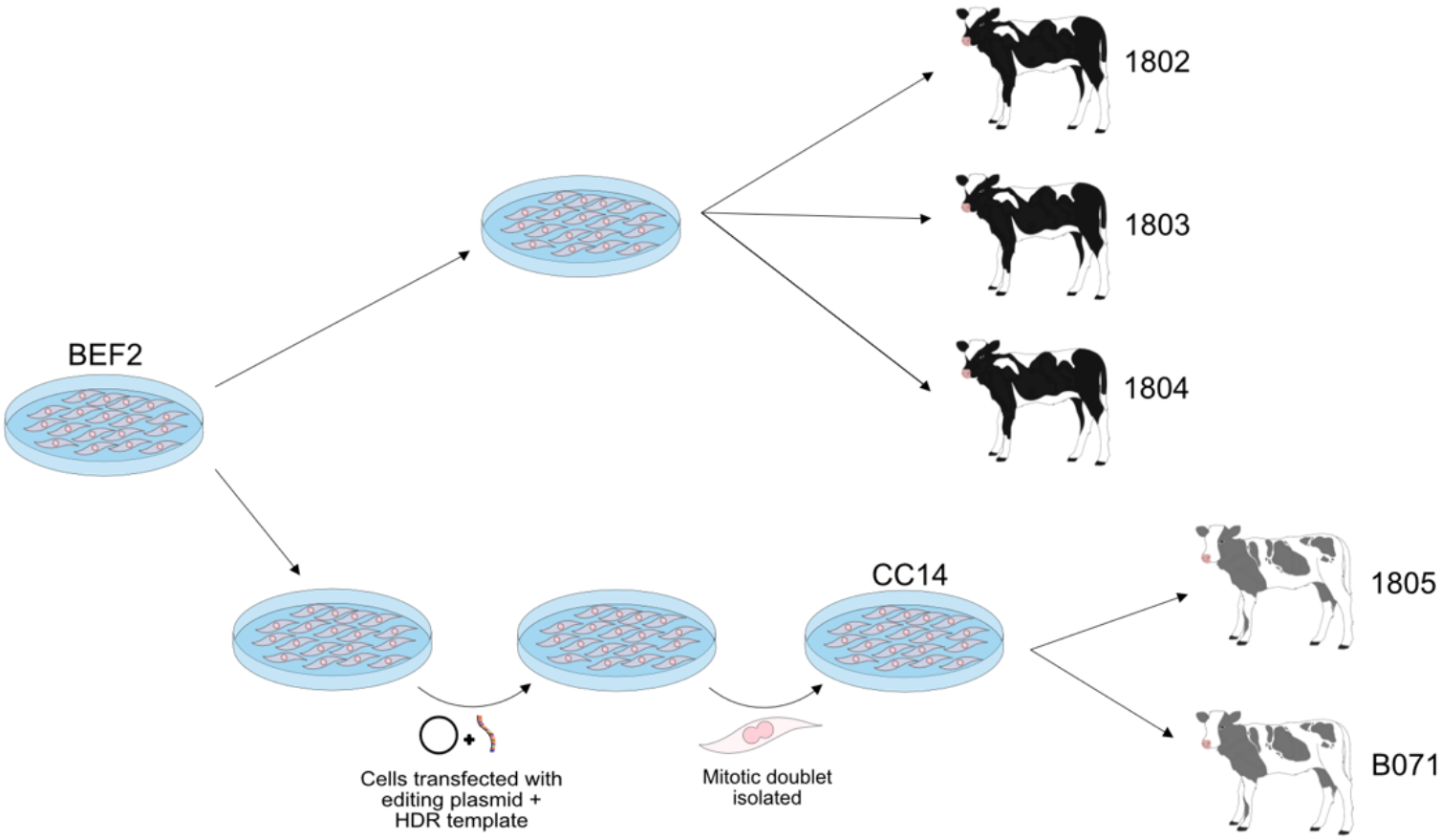
*Relationship between the parental cell line BEF2, edited cell clone CC14 and edited and control calves. Shown is an experimental flow diagram from the parental cell line BEF2 to the two coat colour-diluted Holstein-Friesian calves homozygous for* PMEL *p.Leu18del and three wild-type control calves. A subset of the male primary bovine fetal fibroblasts (BEF2) were transfected with a plasmid-encoded,* PMEL*-specific editor and a single stranded homology directed repair (HDR) template. Post transfection, a single mitotic doublet was used for the clonal isolation of CC14 with a homozygous* PMEL *p.Leu18del mutation. Two edited cloned calves (1805 and B071) and three non-edited control calves (1802, 1803 and 1804) were generated via somatic cell nuclear transfer using CC14 and BEF2 as donor cells, respectively. The ‘named’ samples are those that were sequenced in this study (i.e., BEF2, CC14, 1802, 1803, 1804, 1805, and B071).*

### Identification of off-target mutations from WGS data

To identify mutations that may be the result of CRISPR-Cas9 mediated gene-editing, we applied a series of stringent filtering procedures (Fig 2). Variants relative to the reference genome that were identified to be monomorphic across all samples (n=7,670,567), and those few sites with no coverage in the BEF2 parental cell line (n=14,947), were removed which reduced the 8,021,969 variant sites to 336,455 variants. To remove polymorphic variants that were present in BEF2 but are common to the wider cattle population, all variants that were segregating in a large sequenced New Zealand (NZ) dairy cattle population (see Methods) were also removed, further reducing the number of variants to 31,190. Variants that were present in the gene-edited cell line (CC14) and both gene-edited clones (1805 and B071), but absent in the parental cell line (BEF2) and all three control clones (1802, 1803 and 1804), were retained and variants with a map quality score of less than 60 were removed. This reduced the total number of candidates for variants induced by potential off-target events or spontaneous *de novo* mutagenesis to 457. Variants called to be heterozygous by GATK HaplotypeCaller (22) but identified to have an allele dosage significantly less than 0.5 in the CC14 cell line, 1805, or B071, were defined as candidate mosaic mutations and were filtered out, as it was likely that these mutations occurred after the gene-edited mitotic doublet was isolated (Fig 1). The remaining 218 candidate off-target/*de novo* mutations were then manually examined by visualisation of sequence reads in the Integrative Genomics Viewer (IGV). Using this filtering criteria (Fig 2), we identified 151 candidate mutations that may have resulted from off-target mutagenesis (131 SNVs and 20 indels; Table S1). We also investigated SVs that may have been induced by the use of CRISPR-Cas9. Using a case-control design, Delly (v0.8.1) (23) was used to predict the presence of SVs in CC14, 1805 and B071 that were absent in BEF2, 1802, 1803 and 1804. Using this approach, there were no detectable SVs that were present in the CC14 cell line and the two gene-edited cloned calves, yet absent in all control samples (Table S2).

**Fig 2.**
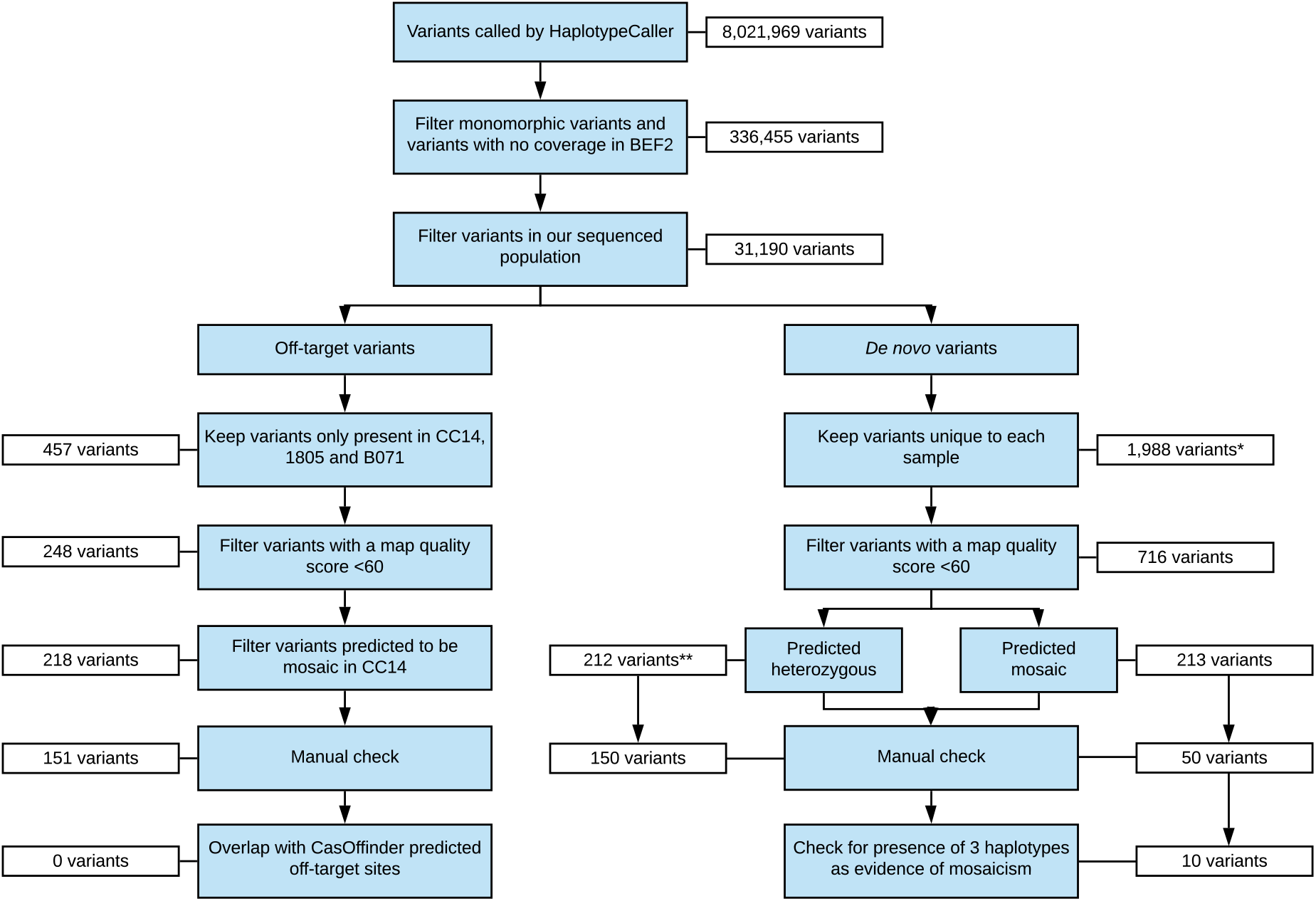
Filtering criteria applied to raw variant calls to identify potential off-target mutations and spontaneous de novo mutations in the gene-edited cell line CC14. The white boxes adjacent the filtering criteria indicate the number of candidate mutations remaining after the filter was applied. *Variants were kept if also present in 1805 and/or B071 **Predicted heterozygous de novo mutations were also filtered for their presence in calves 1805 and B071

### Identification of off-target mutations at predicted candidate loci

Potential genome-wide off-target sites were predicted based on on-target sequence similarity using Cas-OFFinder (12), where we allowed for up to five mismatches with the on-target site. Cas-OFFinder identified 1,166 potential off-target sites, none of which mapped within ±50bp of any of the 151 candidate mutations identified by our discovery pipeline. The sequence flanking each of the 151 candidate mutations was also manually inspected for evidence of sequence similarity with the gRNA and an adjacent PAM site, with no matches or partial matches identified. To ensure that our filtering criteria had not excluded variants from the most likely off-target mutation sites, we also searched the unfiltered variant calls for matches with the sites identified by Cas-OFFinder. We found 230 (of 8,021,969) variants that mapped within 50bp of the 1,166 candidate off-target sites. Almost all (n=225) were filtered out due to being monomorphic across all samples, three sites were filtered out due to poor read quality, one captured the on-target mutation at the edited site, and one site was called in the sample of the non-edited control calf 1803. These steps provided reassurance as to the filtering criteria applied, and suggested that if CRISPR-Cas9 induced off-target mutagenesis had occurred, it had not done so at any of the most biologically plausible sites.

### Long molecule sequencing of the on-target site

To investigate the on-target edit site for SVs and plasmid integration events, we conducted long-range polymerase chain reaction (PCR) to amplify approximately 8.8kb of sequence surrounding the edit site (BTA5:57,340,856bp-57,349,715bp) in the parental cell line (BEF2), the gene-edited cell clone (CC14), two gene-edited cloned calves (1805 and B071), and one control clone calf (1802). The amplicons were sequenced using the Oxford minION platform, generating an average sequence depth of 590x across the targeted region in each of the five samples, and minimap2 (24) was used to map the long sequence reads to the bovine ARS-UCD1.2 reference genome (21). Since structural variation might disrupt primer binding and lead to allele drop-out at the locus (i.e., a large hemizygous structural variant that could confound PCR), we looked for collocating variants to confirm biallelic amplification of the region. Manual inspection of the sequence reads in IGV revealed two such biallelic SNVs (BTA5:57,343,664G>A and BTA5:57,348,336G>A) heterozygous in these samples, confirming that we captured both the maternal and paternal haplotypes across this region. Alignment of the long reads to the *PMEL*-specific CRISPR-Cas9 expression plasmid sequence using minimap2 (24) revealed no matches, suggesting that the editing plasmid was unlikely to have integrated at the on-target site.

### Investigating evidence of plasmid integration

Although a PCR assay had previously failed to amplify a specific plasmid fragment (20), that approach assumes contiguous sequence representation of the plasmid template, and thus WGS data allows a more comprehensive analysis of potential integrations of the *PMEL*-specific CRISPR-Cas9 expression plasmid (and any potential fragments thereof). To investigate possible plasmid integration events at sites other than the on-target site, we added the sequence of the *PMEL*-specific CRISPR-Cas9 expression plasmid, that had been used for editing (a pX330 derivative), to the ARS-UCD1.2 reference genome (21) and re-ran sequence alignments using the Burrows-Wheeler Aligner (BWA; (25)) for the parental cell line (BEF2), gene-edited cell line (CC14), two gene-edited cloned calves (1805 and B071), and one control clone calf (1802). In all samples we observed a pile up of sequence reads in a G-rich repeat region at 828-873bp on the *PMEL*-specific editing plasmid. The mapping quality scores ranged between 0-35, suggesting these were mismapped reads, and of reduced interest given these were not polymorphic across the control and edited samples. No additional sequence reads were observed to map to the plasmid sequence for the two edited calves, control calf and parental cell line. Only for the CC14 sample, we found 46 additional sequence reads that appeared to map to the plasmid sequence (maximum coverage of 8x). A *de novo* assembly of these reads indicated that these reads could not be assembled into a single contiguous sequence, and alignment to the bovine genome using BLAST (26) did not highlight any sequence overlap.

The limited read representation of *PMEL*-specific editing plasmid sequences mapped in CC14, and lack of these sequences in CC14-derived animals suggested bi- or mono-allelic integration in CC14 was unlikely, however we performed additional experiments to further investigate this possibility. Here, we designed two PCR primer pairs that together covered 1,365bp of the plasmid region, targeting sequence that overlapped the regions of homology identified from the short-read alignments. We designed a single primer pair targeted at BTA2:110,817,757-110,818,275bp, representing *Bos taurus* genomic sequence that would be expected to amplify in all samples. We created a mock plasmid-integrated DNA sample by spiking in 0.14pg of the *PMEL*-specific editing plasmid into BEF2 gDNA, aiming to simulate a sample with a single integration event and thereby act as a positive control. These PCRs were conducted on DNA extracted from BEF2, B071, 1805, 1802, an aliquot of CC14 DNA previously extracted for WGS, and a fresh sample of DNA extracted from the CC14 cell clone. PCR amplification of the plasmid-specific 757bp and 690bp fragments returned a positive result in the plasmid and positive control sample, but a negative result in BEF2, both CC14 samples, B071, 1805 and 1802 (Fig 3). These results suggest that the short-read data seen to map to the plasmid sequence in CC14, was unlikely indicative of an integration event, and more likely due to low levels of sample contamination prior to WGS.

**Fig 3:**
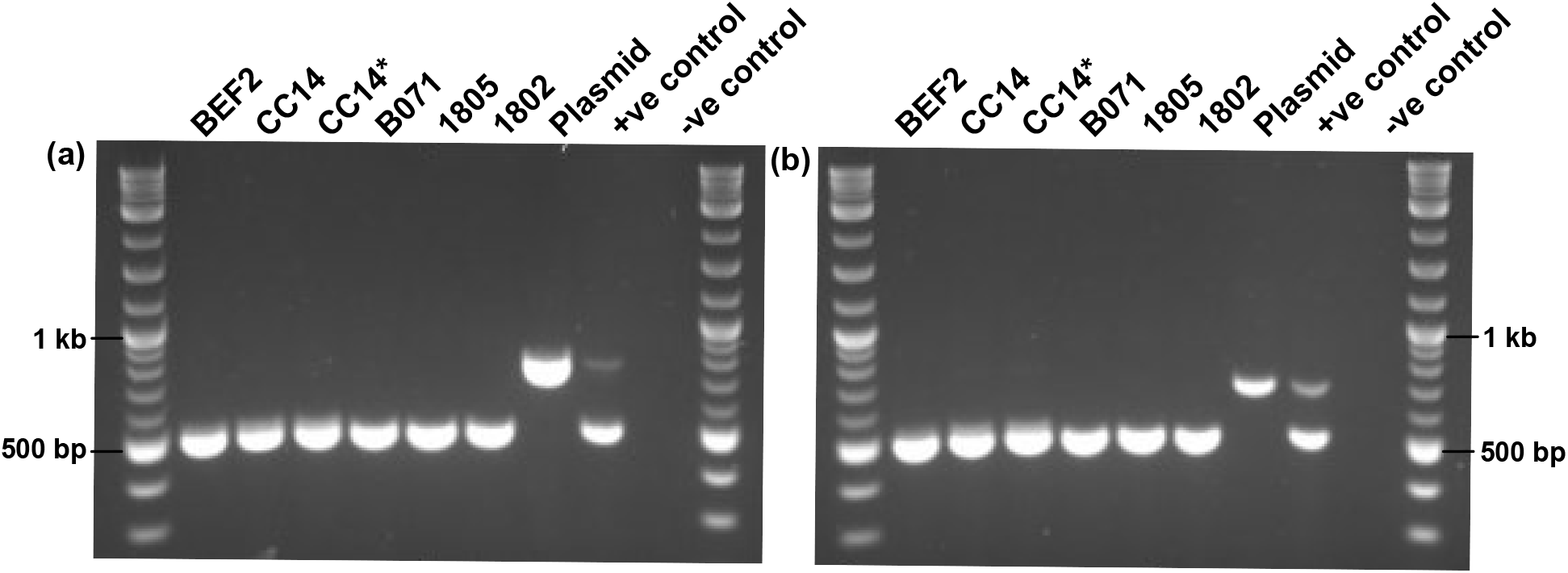
Absence of editing plasmid-specific fragments in genomic DNA extracted from the parental cell line (BEF2), the gene-edited cell clone CC14, DNA sent away for WGS of CC14 (CC14*), and genomic DNA extracted from cloned calves B071, 1805, and 1802. Each PCR reaction contained two sets of primers and BEF2 spiked in with 0.14pg of plasmid DNA was used as the positive control. (a) Primer pair designed to amplify bovine BTA2:110,817,757-110,818,275bp (519bp), and another designed to amplify CRISPR-Cas9 expression plasmid-specific region 6,263-7,019bp (757bp); (b) Primer pair designed to amplify bovine BTA2:110,817,757-110,818,275bp (519bp product), and another to amplify plasmid-specific region 6,939-7,628bp (690bp).

**Fig 3.**
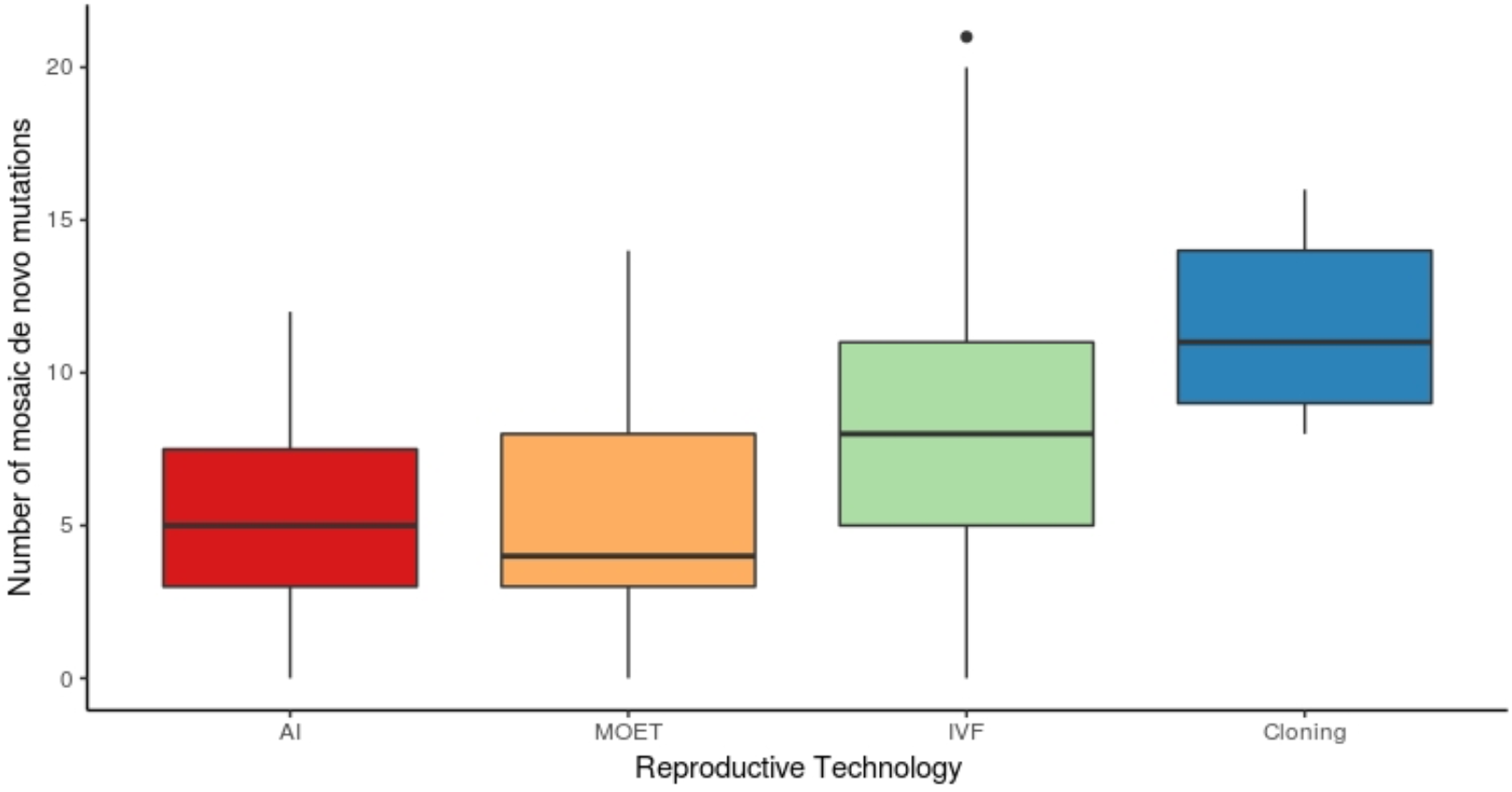
Comparison of the number of mosaic de novo mutations observed in cattle born from the use of artificial insemination (AI), multiple ovulation embryo transfer (MOET), in vitro fertilisation (IVF) and cloning.

### Analysis of de novo mutations in the cloned calves

The cloned calves used for this study were generated by somatic cell nuclear transfer with donor cells from either the parental cell line BEF2, or the gene-edited cell clone CC14 (20). To identify *de novo* mutations carried by each cloned calf, either originating from the donor cell or occurring during development of the calf, we applied the filtering criteria outlined in Fig 2. To differentiate between *de novo* mutations that likely occurred in cell culture and were subsequently inherited by the cloned calves, from *de novo* mutations that likely occurred during development of the cloned calves (i.e., after first cell division), we categorised *de novo* mutations as heterozygous or mosaic based on allele dosage at each site (Table 1). A binomial probability function was applied to determine if the allele dosage at each variant site was consistent with a truly heterozygous genotype expected for a *de novo* mutation already present in the donor cell. When the allele dosage at a variant site was determined to be not statistically different from the expected allele dosage of 0.5, the variant was predicted to be a candidate heterozygous *de novo* mutation in the cloned calf, whereas if allele dosage was significantly less than 0.5, the variant was predicted to be a candidate mosaic *de novo* mutation that arose during development of the cloned calf. All variants were manually assessed in IGV software, after which a proportion of candidate *de novo* mutations were filtered out due to representing incorrect variant calls, most often due to errors based on proximity to polynucleotide regions, repetitive regions, miscalled variants in other samples, proximity to indels, or misalignment of reads. Table 1 shows the number of variants that remained after applying the filtering criteria outlined under ‘*de novo* variants’ in Fig 2, where ‘likely *de novo*’ mutations are those that remained after the manual check.

**Table 1.** Number of candidate de novo mutations identified after each filter was applied to 31,190 filtered variants across the three control cloned calves, two gene-edited cloned calves and gene-edited cell clone

| Sample ID | Unique to each sample | Map quality = 60 | <u>Heterozygous *de novo* mutations</u> |  | <u>Mosaic *de novo* mutations</u> |  |
| --- | --- | --- | --- | --- | --- | --- |
|  |  |  | Candidate *de novo* | Likely *de novo* | Candidate *de novo* | Likely *de novo* |
| 1802 | 1,224 | 439 | 340 | 205 | 45 | 16 |
| 1803 | 1,402 | 550 | 409 | 276 | 82 | 14 |
| 1804 | 2,769 | 686 | 566 | 293 | 67 | 9 |
| 1805 | 1,145 | 433 | 313 | 197 | 52 | 8 |
| B071 | 1,470 | 587 | 408 | 277 | 133 | 11 |
| CC14 | 1,988 | 716 | 212* | 150 | 213 | 50 |

#### Heterozygous de novo mutations

The majority of *de novo* mutations present in the cloned calves appear to be heterozygous variants and are likely inherited from the donor cell used for somatic cell nuclear transfer. A pairwise comparison of the number of likely heterozygous *de novo* mutations inherited by each of the cloned calves (Table 1) suggests that the number of mutations observed in each clone is statistically different between six of the ten pairs (Table 2). The pair-wise comparison does not draw a distinction between the number of heterozygous mutations observed in the gene-edited compared to the non-edited calves, but rather appeared random. Based on these results, the number of heterozygous *de novo* mutations inherited by cloned calves generated from the gene-edited cell clone CC14 (1805 and B071) did not appear to be different than those in cloned calves generated from the non-edited, parental cell line BEF2 (1802, 1803 and 1804).

**Table 2:** Results (p-values) from two-proportion z-test comparing the difference in number of likely heterozygous (top) and mosaic (bottom) de novo mutations observed in the cloned calves.

|  | 1802 | 1803 | 1804 | 1805 | B071 |
| --- | --- | --- | --- | --- | --- |
| 1802 | | $1.36 \times 10^{-3}$ | $9.07 \times 10^{-5}$ | 0.73 | $1.17 \times 10^{-3}$ |
| 1803 | 0.86 | | 0.5 | $3.18 \times 10^{-4}$ | 1 |
| 1804 | 0.23 | 0.4 | | $1.64 \times 10^{-5}$ | 0.53 |
| 1805 | 0.15 | 0.29 | 1 | | $2.7 \times 10^{-4}$ |
| B071 | 0.44 | 0.69 | 0.82 | 0.65 |  |

#### Mosaic de novo mutations

The number of mosaic *de novo* mutations identified was more than a magnitude lower than the number of heterozygous *de novo* mutations identified (Table 1). These mutations, occurring during development of a calf, would be expected to be in complete, but imperfect linkage with the paternal or maternal haplotype (27), and we would therefore expect to see three haplotypes at the variant site. Each ‘likely *de novo*’ mosaic mutation (Table 1) was manually checked for evidence of a segregating bi-allelic variant on the same read, or read pair, to support the presence of three haplotypes and strengthen the evidence supporting a true mosaic mutation. Out of the total number of variants predicted to be likely true mosaic mutations: 8/16 variants in 1802, 6/14 variants in 1803, 5/9 variants in 1804, 2/8 variants in 1805, and 5/11 variants in B071 had evidence of three haplotypes and could be confirmed as true mosaic *de novo* mutations. A pair-wise significance test demonstrated that the difference in number of likely mosaic *de novo* mutations carried by each cloned calf (Table 1) does not appear to be statistically significant, regardless of the cell line of origin (smallest *p-*value = 0.15 between calves 1802 and 1805; Table 2). These results suggest that the *de novo* mutation rate during embryonic development does not significantly differ between cloned calves generated using donor cells from a cell clone edited by the CRISPR-Cas9 gene-editing tool, and those generated using a non-edited cell line of the same parental origin.

#### Comparison of mosaic de novo mutation rate in cloning compared to other reproductive technologies

Since we were unaware of any study to date that has attempted to quantify the *de novo* mutation rate in cloned animals, we compared the number of mosaic *de novo* mutations that occurred in the cloned calves described in this study, with the number of mosaic *de novo* mutations reported for generation of animals using other reproductive technologies. Here, the rates of mosaic *de novo* mutation for our clones (n=5) were contrasted with those observed in cattle generated via artificial insemination (AI; n=35), multiple ovulation embryo transfer (MOET; n=44), and in vitro fertilisation (IVF; n=43), where these other data were derived from 131 three or four generation pedigrees previously published by Harland et al. (28). Acknowledging the comparatively smaller sample size of our study, these results suggest that the mosaic *de novo* mutation rate in cloned calves is significantly higher than what is observed with the application of AI (*p*=0.0097) and MOET (*p*=0.012), but not significantly higher than that observed with IVF (*p*=0.13).

### Comparison of *de novo* mutation distribution and spectra

To further evaluate the candidate *de novo* mutations across experimental conditions, we categorised mutations according to the different stage of their occurrence, and compared mutation distribution and spectra of mutations occurring at each of these stages. *De novo* mutations that arose in cells post plasmid transfection (n=150; Table 1) were estimated based on heterozygous *de novo* mutations in the CC14 cell clone that were subsequently inherited by cloned calves B071 and 1805, but absent in all other samples. The number of *de novo* mutations that emerged during cell clone expansion were estimated based on the sum of mosaic *de novo* mutations in the gene-edited cell clone CC14, and heterozygous mutations that were present in any cloned calf, but not in CC14 or the parental cell line, BEF2 (n=1298; Table 1). The smallest proportion of *de novo* mutations (n=58; Table 1) arose during the development of the cloned calves. Across the three groups, *de novo* mutations appeared to be randomly dispersed across most of the genome and were not observed to cluster in a group-dependent manner (Fig 4a), but a distinct spectra of mutations was observed between *de novo* mutations that were predicted to have arisen in the cloned calves and those that were predicted to have arisen post plasmid transfection or during cell culture (Fig 4b). Comparison of mutation spectra between the three groups revealed that C>A mutations were significantly enriched in cells post plasmid transfection and cells in culture for clonal expansion, compared to those in the cloned calves (*p*=3.213×10^−6^), accounting for over 40% of total mutations observed in cells at the two stages of *in vitro* cell culture. The cloned calves appeared to be significantly enriched for T>G mutations compared to cells post plasmid transfection and cells in culture (*p*=3.213×10^−6^). These mutations accounted for 31% of the total *de novo* mutations observed in the cloned calves. There were no significant differences in mutation spectra observed between cells post plasmid transfection and cells in culture.

**Fig 4.**
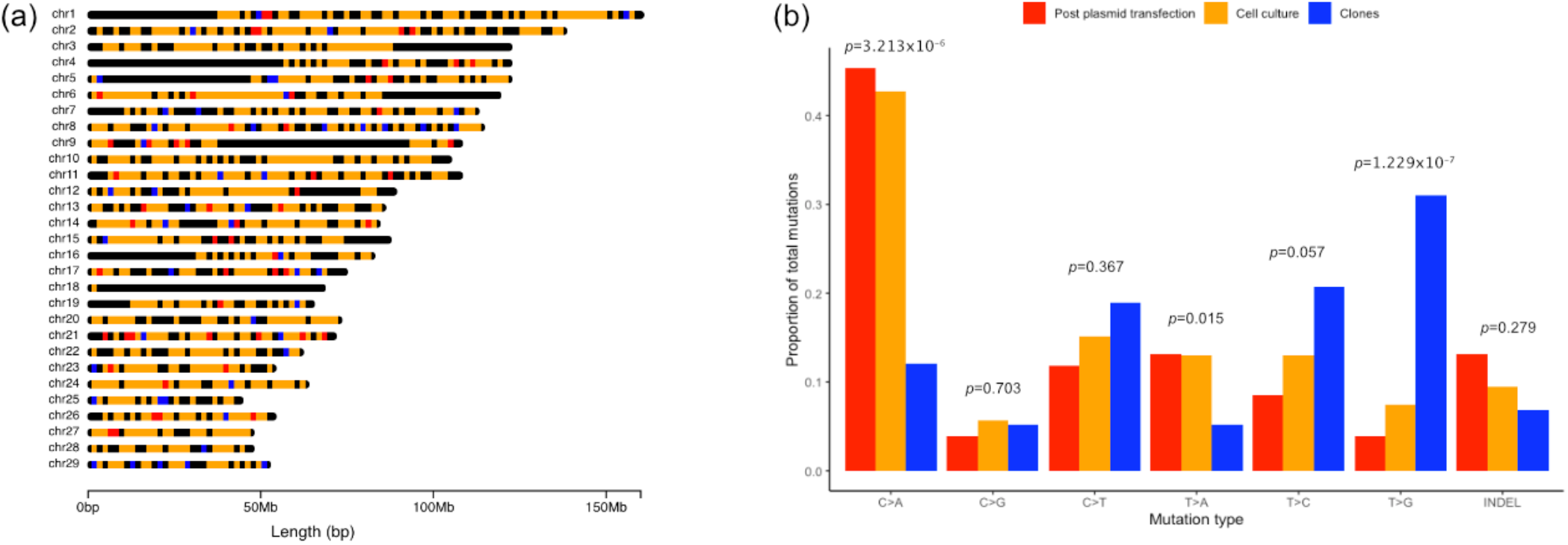
Distribution and spectra of de novo mutations predicted to have arisen in cells post plasmid transfection (red), during the cell culture expansion phase (orange) and during development of the cloned calves (blue). (a) Distribution of de novo mutations (SNVs and indels) across the bovine genome. (b) Proportion of de novo mutations within each mutation class observed between groups. A fisher’s exact test comparing the proportion of observed mutations per mutation class between groups, showed a significant difference in C>A mutations (p=3.213×10^−6^) and T>G mutations (p=1.229×10^−7^).

## Discussion

We present the first study in cattle based on cloned calves produced by somatic cell nuclear transfer to evaluate unintended off-target mutations, SVs at the on-target site, and unintended integration of the editing plasmid associated with the application of CRISPR-Cas9 mediated gene-editing. Using WGS data and long molecule sequencing, we show that the application of CRISPR-Cas9 to induce a precise 3bp deletion in the *PMEL* gene did not produce detectable off-target events in the gene-edited cell clone (CC14) or the resultant gene-edited calves (1805 and B071). Furthermore, we provide primary evidence to suggest that CRISPR-Cas9 mediated gene-editing does not affect spontaneous mutagenesis or mutation spectra in subsequent cell divisions post-edit, and *de novo* mutagenesis in calves derived from the gene-edited cell clone appears to be equivalent to that of controls.

The filtering criteria that we used in this study was built around our clone-based study design where each sample originated from the same genetic background. The study design, combined with high-depth WGS data enabled direct comparisons between spontaneous mutagenesis that occurred in cells post plasmid transfection, in cell culture during cell clone expansion, and during development of the cloned calves. Application of these filtering criteria combined with manual sequence visualisation revealed no detectable CRISPR-Cas9 induced off-target SNVs, indels or SVs in the gene-edited cell clone or gene-edited cloned calves. Although integration events for circular, supercoiled plasmids are rare (29), it was possible that the whole plasmid, or parts of the editing plasmid, may have integrated at the on-target site or elsewhere in the genome (10,30,31). For this reason, we targeted a broad 8.8kb interval at the on-target site for high-depth long molecule sequencing. Although the results from this did not reveal evidence of an integration event or other structural variation, this did not rule out the possibility of whole or partial integration of the vector at an off-target site. To investigate this possibility, we added the *PMEL*-specific editing plasmid sequence to the reference genome and re-ran our short-read sequence alignment. Several reads from the CC14 gene-edited cell clone were found to map to this sequence, but our follow-up PCR analysis showed that these reads were most likely the result of sample contamination prior to WGS, rather than evidence of an integration event. While somewhat surprising, residual contamination events have been previously reported in other sequencing contexts (32), and so it seems plausible that the contamination noted here may have occurred during one of the many handling steps prior to WGS. Our findings highlight the importance of a methodical approach to investigating plasmid integration events when using double-stranded DNA to deliver editing tools such as CRISPR-Cas9, and the need for carefully designed experiments to ensure fragments of the plasmid do not persist in the edited genome. Delivering editors as purified proteins (i.e., ribonucleoprotein complexes) could be assumed to minimise this risk and as such represent an appealing alternative to plasmid-based methods.

The number of heterozygous *de novo* mutations varied significantly between calves, but these changes could not be attributed to any single experimental condition (i.e., cell line, gene editing status, or donor cell origin). As we expect heterozygous *de novo* mutations in the cloned calves to have been inherited from their somatic donor cells, these results suggest that the use of CRISPR-Cas9 gene-editing is unlikely affecting the expected spontaneous mutation rate during clonal expansion of the gene-edited cell. The difference in observed heterozygous *de novo* mutations may instead be due to differences in the accumulation of unrepaired DNA damage across cells in culture, which could be induced by oxidative damage, damage due to UV exposure, other mutagens, or mechanical sheer (33,34). Indeed, when we examined the mutation spectra for these mutations, we observed a significant enrichment in C>A mutations, a base-pair transversion associated with cells in culture and thought to be caused by oxidative stress (35,36). By contrast, the mosaic *de novo* mutations observed in the cloned calves appeared to be statistically equivalent regardless of their edited or non-edited status, although it is important to note that some somatic mutations may not be represented in the WGS data or discarded as sequencing errors due to their low-abundance or absence in the sampled tissue. When the frequency of *de novo* mutations was compared to that observed in cattle generated with the assistance of reproductive technologies such as AI, MOET and IVF (28), we observed that the average number of *de novo* mutations reported for the cloned calves was greater than the average number observed in the other groups, but not significantly so when compared to IVF. The increased rate of mutagenesis observed in cloning and IVF compared to other reproductive technologies may be due to potentially suboptimal *in vitro* culture conditions and manipulations that are at the center of these technologies. Analysis of the *de novo* mutation spectra revealed a marked difference in predominant mutation type between the cloned calves and cattle produced by natural matings and other reproductive technologies (37). We observed a significant enrichment in T>G transversion mutations in the cloned calves, where an excess of C>T transition mutations is usually seen in cattle born from a natural mating or other assisted reproductive technologies. The T>G mutation type has been observed to be enriched amongst mouse somatic mutations and thought to be attributable to less effective repair of thymine dimers, but the exact mechanism of mutagenesis remains unconfirmed (38), and why this transversion is enriched in the cloned calves remains unclear. The results presented here must be interpreted with caution due to the small sample size used for comparison, and a larger dataset will be required to support these findings.

Our results demonstrate that naturally occurring, beneficial genetic variation can be introduced into animals that subsequently show levels of mutagenesis indiscernible from the *de novo* mutation rates of un-edited controls. Although gene-editing technologies such as CRISPR-Cas9 hold potential to accelerate introgression of favourable genetic variants across large populations, the widespread use of such technologies is limited in the agricultural sector due to uncertainties around the level of social acceptance and controversy surrounding perceived safety of the products from gene-edited animals. A major challenge is the ability to detect and quantify off-target mutagenesis above the background *de novo* mutation rate. We can identify candidate off-target events using logical filtering criteria and evaluate each site for its biological plausibility based on sequence similarity with the on-target site, but it is more difficult to differentiate between off-target mutations and spontaneous *de novo* mutations at regions with little homology to the on-target site. Holstein-Friesian cattle have a baseline spontaneous *de novo* mutation rate of approximately 1.2×10^−8^ mutations per bp, per generation (28).

Quantification of off-target mutations at sites of little homology would therefore require tens, potentially hundreds, of extra mutations in edited samples compared to controls, to observe a ‘significant’ increase in mutation rate above baseline levels. Spontaneous *de novo* mutations have been observed to follow a typical signature, where there is an expected excess of C>T mutations (27). Comparing mutation spectra between edited and non-edited samples may thus prove useful in evaluating the occurrence of unintended mutagenesis, although we are unaware of any studies that have specifically investigated the mutation profile of off-target events induced by gene-editing technologies. Development of sensitive tools that enable accurate detection of genuine off-target events, but also consider natural *de novo* mutation, may be difficult but will aid to establish the risk profile of gene-editing technologies and ultimately support informed consumer decisions.

## Methods and Materials

### Animal generation

All animals and cell lines described in the present study were generated as reported by Laible et al. (20). Briefly, male primary bovine fetal fibroblast cells (BEF2) were co-transfected with a modified pX330 transfection vector carrying Cas9 nuclease and *PMEL*-specific gRNA, and a homology-directed repair template using a Neon transfection system (Invitrogen). Two days post transfection, mitotic doublets were manually selected, reseeded and cultured. Cell clones identified to be homozygous for the targeted 3bp deletion in exon 1 of the *PMEL* gene (p.Leu18del; BTA5:57,345,301-57,345,303bp) were further expanded (CC14). Donor cells from the biallelic cell clone CC14 and the wildtype parental cell line BEF2 were used to generate two *PMEL* gene-edited calves (1805 and B071) and three wildtype calves (1802, 1803 and 1804), respectively, via somatic cell nuclear transfer.

### Whole genome sequencing and data analysis

Unedited male primary bovine fetal fibroblast cells (BEF2), edited fetal fibroblast cells homozygous for p.Leu18del (CC14), three control clone calves generated from BEF2, and two gene-edited clone calves generated from CC14 were chosen for whole genome sequencing. Genomic DNA was isolated from CC14 and BEF2 cells, and blood samples from each of the calves using a Nucleon BACC2 kit (Cytiva, Little Chalfont, UK). The genomic DNA samples were sequenced by Macrogen (Seoul, South Korea), with a targeted read depth of 60x per isolate. The samples were sequenced based on 150bp paired reads on the Illumina HiSeq X Ten platform and read mapping was performed using the ARS-UCD1.2 genome build (21) and the BWA MEM v0.7.17 software (39), resulting in mean mapped read depth of 50.7x across the genome (ranging between 44.7x to 54.8x across samples). SNV and indel calling was carried out using Genome Analysis Toolkit (GATK) HaplotypeCaller (v4.0.2.1) using default parameters (22), yielding an unfiltered dataset of 8,021,969 variants across the seven samples.

### Identification of off-target mutations

All 8,021,969 variants called by HaplotypeCaller (v4.0.2.1) (22), were dummy coded (0 = no coverage; 1 = homozygous reference; 2= heterozygous; 3 = homozygous alternate). Variants identified to be monomorphic across all samples were removed, sites with no coverage in BEF2 were removed, and all variants present in an unrelated sequenced cattle population previously described by Jivanji et al. (40), and Lopdell et al. (41) (n = 564, remapped to the ARS-UCD1.2 genome build (21) yielding 37,208,259 SNPs and 11,746,534 indels) were removed. Candidate off-target mutations were filtered according to the following criteria: (1) candidate mutations should be present in the CC14 cell clone and in both gene-edited clones, but absent in the BEF2 cell clone and three control clones, (2) should have a map quality score of 60, (3) should have an allele dosage of, or statistically equivalent to, 0.5 or 1 for the alternative allele in the CC14 cell clone and both gene-edited clones, and (4) manual inspection of sequence reads should show no evidence of miscalled or misaligned SNVs/indels at the candidate positions. Allele dosage was calculated for each variant by dividing the number of alternate reads by the total number of observed reads at each position. A binomial probability function was used to predict if the allele dosage was statistically equivalent to 0.5 for a heterozygous genotype, with a Bonferroni corrected *p*-value calculated as the significance threshold. In practice, these criteria would highlight a 60x depth site as being a potentially mosaic variant with a 10:50 depth ratio. All candidate off-target mutations were uploaded into IGV for visualisation (42), and the sequence adjacent each candidate off-target mutation was visually inspected for sequence similarity with the gRNA (ATGGGTGTTCTTCTGGCTGT) and the presence of a 5’-NGG-3’ PAM site.

Potential off-target sites were also predicted using Cas-OFFinder software (12). The online Cas-OFFinder tool was used to identify potential off-target mutations by searching the ARS-UCD1.2 genome build (21) for sequence similarity with the gRNA used to target the *PMEL* gene, allowing for up to five mismatches. Candidate off-target mutations predicted by the software were compared to a list of candidate off-target mutations identified by the filtering criteria described above, and also to the unfiltered variants called by GATK HaplotypeCaller (v4.0.2.1) (22).

Candidate SVs that may have arisen due to the application of CRISPR-Cas9 gene-editing were called and filtered using Delly (v0.8.1) (23). A case-control approach was implemented in Delly where the CC14 cell clone, clone 1805 and clone B071 were separately called as case samples with the parental cell clone, BEF2, used as the control. After initial SV calling, the wild type clones generated from BEF2 (1802, 1803 and 1804), were added as additional controls to further filter candidate SVs. All candidate mutations were manually inspected in IGV (42) to assess evidence of a legitimate SV at each of these sites.

### Long molecule sequencing

Genomic DNA was extracted from cultured bovine cells for samples BEF2 and CC14, and from blood samples for 1805, B071 and 1802 as previously described by Laible et al. (20). Primers were designed to target BTA5:57,340,856-57,349,715bp (Table S3), encapsulating 8,860bp around the *PMEL* on-target site. The PCR was conducted using the KAPA LongRange PCR kit (KapaBiosystems) with the following cycling conditions: 95°C for 3 minutes; 95°C for 30 seconds, 60°C for 30 seconds, and 68°C for 9 minutes for 35 cycles; and a final extension step of 68°C for 9 minutes. The PCR products were loaded and run on a 1% agarose gel for 60 minutes at 100 V to estimate amplicon size. Resultant amplicons were purified using AMPure XP beads and then used to construct a sequencing library using the SQL-LSK109 kit (Oxford Nanopore Technologies) as per the manufacturer’s instructions. The library was constructed using 700ng of DNA from across the five samples, loaded onto a FLO-MIN106 flow-cell (Oxford Nanopore Technologies) and sequenced for 10 minutes, yielding an average 590x coverage over BTA5:57,340,856bp-57,349,715bp for each sample. The reads were base-called using Guppy basecaller (v4.0.14) (43), with the samples then separated based on their barcodes by Guppy barcoder (v4.0.14), and subsequently aligned to the ARS-UCD1.2 reference genome (21) plus the *PMEL*-specific CRISPR-Cas9 expression plasmid sequence using minimap2 (v2.14) (24).

### Investigation of the presence of the PMEL-specific CRISPR-Cas9 expression plasmid

The genomic DNA samples described above were also used for PCR. Two primer pairs were designed across the *PMEL*-specific CRISPR-Cas9 expression plasmid sequence, where targeted regions were chosen based on mapped short read WGS data from the CC14 cell clone (Table S3). Each PCR reaction contained two sets of primer pairs at a concentration of 5μM per primer: one primer pair specific for the editing plasmid, and another primer pair targeted to amplify BTA2:110,817,757-110,818,275 (Table S3). The PCR was conducted using the Kapa 2G Fast Hotstart PCR kit (KapaBiosystems) with the following cycling conditions: initial denaturation at 95°C for 3 minutes; denaturation at 95°C for 15 seconds, anneal at 60°C for 15 seconds, extend at 72°C for 15 seconds, for a total of 35 cycles; and final extension at 72°C for 1 minute.

### Identification of *de novo* mutations

*De novo* SNVs and indels unique to each sample were identified using a filtering criteria similar to that described above for identifying off-target mutations. As described above, variants were initially filtered for monomorphic sites, sites missing in BEF2 and alleles already identified to segregate in a sequenced NZ dairy cattle population. From the remaining variants, *de novo* mutations were identified by the following criteria: (1) keeping heterozygous SNVs and indels specific to each sample; (2) filtering to remove reads with a map quality score less than 60; (3) classifying SNVs and indels as heterozygous or mosaic *de novo* mutations, where mosaic variants were defined as having an allele dosage significantly less than 0.5, as determined by the binomial probability function described previously; (4) filtering variants based on manual examination of sequence alignments in IGV to remove misaligned or miscalled SNVs and indels. Sequence alignments were also examined at each candidate mosaic SNVs and indel site for evidence of more than one bi-allelic variant segregating on the sequence read, or read pair, that could indicate the presence of three haplotypes and support mosaicism. Pair-wise comparisons of *de novo* mutation rates in the cloned calves were conducted using a two-proportions Z-test, and comparisons of *de novo* mutation spectra were conducted using Fisher’s exact test.

## Acknowledgments

The authors would like to acknowledge Massey University and Livestock Improvement Corporation (LIC) for their support in this research, AgResearch for the generation of this dataset, The University of Auckland for access to laboratory resources, and the New Zealand eScience Infrastructure (NeSI) for providing the computational resources required for the analyses described here. This work was funded by AgResearch and the Ministry of Business, Innovation and Employment.

## Supporting information

### Additional File 1 Table S1

Format: .csv

Title: Predicted and candidate off-target mutations.

Description: All predicted and candidate off-target mutations identified by our filtering criteria, with additional information about mutation type, genes that the mutations may map within, and the predicted variant effect.

### Additional File 2 Table S2

Format: .dox

Title: Structural variants (SVs) identified in the gene-edited cell line (CC14) and gene-edited cloned calves (1805 and B071) using DELLY with the parental cell line (BEF2) and non-edited cloned calves (1802, 1803 and 1804) as reference samples.

### Additional File 2 Table S3

Format: .dox

Title: Description of PCR primer pairs designed to investigate the on-target site and plasmid integration.

## Notes

### Competing Interest Statement

ML and CH are employees of Livestock Improvement Corporation, a commercial provider of bovine germplasm. The remaining authors declare that they have no competing interests.

