## Additional File 2 for "The genomes of precision edited cloned calves show no evidence for off-target events or increased *de novo* mutagenesis"

Table S2 Structural variants (SVs) identified in the gene-edited cell line (CC14) and gene-edited cloned calves (1805 and B071) using DELLY with the parental cell line (BEF2) and non-edited cloned calves (1802, 1803 and 1804) as reference samples

|  | **CC14** | **1805** | **B071** |
| --- | --- | --- | --- |
| BEF2 as reference sample | 27 | 38 | 32 |
| 1802, 1803 and 1804 added as reference samples | 1 | 5 | 5 |
| SVs common between gene-edited samples | 0 | 0 | 0 |

Table S3 Description of PCR primer pairs designed to investigate the on-target site and plasmid integration

|  | Sequence | Melting Temp ( ֯C ) | PCR product size | Position |
| --- | --- | --- | --- | --- |
| **Long-range PCR primers** | | | | |
| Forward primer | GTGCCACTGACATGTAGCAAAG | 60.8 | 8,860bp | BTA5:57,340,856-57,349,715bp |
| Reverse primer | CCCTCCTCAGTCCTTACCAGTA | 59.6 |  |  |
| **Vector integration PCR primers** | | | | |
| **Set 1** |  |  |  |  |
| Forward primer | TGACGTTGGAGTCCACGTTC | 62.1 | 757bp | gRNA/Cas9 plasmid:6,263-7,019bp |
| Reverse primer | TCTTCGGGGCGAAAACTCTC | 63.9 |  |  |
| **Set 2** |  |  |  |  |
| Forward primer | AGATCAGTTGGGTGCACGAG | 61.3 | 690bp | gRNA/Cas9 plasmid:6,939-7,628bp |
| Reverse primer | TGACTCCCCGTCGTGTAGAT | 60.5 |  |  |
| **Internal control PCR primers** | | | | |
| Forward primer | ATGTTAGGTGCAGGTGGAGC | 60.1 | 519bp | BTA2:110,817,757-110,818,275bp |
| Reverse primer | GCTTCCCACCTTGACCTCTC | 61.2 |  |  |
